# Dissecting the metabolic costs of up- and down-hill walking

**DOI:** 10.1101/2024.08.08.607266

**Authors:** Luke N. Jessup, Luke A. Kelly, Andrew G. Cresswell, Glen A. Lichtwark

## Abstract

Work- and collision-based models of locomotion are often used to describe the relationship between the body’s mechanical work requirements and metabolic energy expenditure. While work- and collision-based models do a reasonable job of relating mechanical work to metabolic cost at a system level, these models may not map to the underlying force and work demands of muscle, which directly affects energy expenditure. We collected motion capture, force, electromyography and ultrasound data from the main power producing muscles during uphill and downhill walking between +/− 15% grade. These data were used to evaluate a musculoskeletal modelling approach to simulate muscle force- and work-related costs that could be compared to metabolic power that we measured using indirect calorimetry. Muscle force-related costs (activation heat rate + maintenance heat rate) increased at steeper up- and down-hill grades and were moderately correlated with mean joint moments. Muscle work-related costs (mechanical work rate + shortening / lengthening heat rate) increased as grade became more positive and were strongly correlated with net joint work. Compared to traditional models, the inclusion of a term to account for muscle force-related costs should lead to a more explanatory cost model that maps directly to the mechanical demands of muscle.

## Introduction

Locomotion is rarely performed at steady state or on level ground. Our body’s mechanical energy (work) requirements are continually fluctuating. We can anticipate increases or decreases in these work requirements, but the precise demand placed on the musculoskeletal system, including the metabolic costs, are not well understood. This information is necessary to form a cohesive biomechanical understanding of how energy is added to, or absorbed by the system whenever we locomote up- or down-hill, on more or less compliant surfaces, with more or less mass, accelerating or braking etc. This study focuses on graded locomotion, where changes in surface gradient result in proportionate change in mechanical work requirements, making it an ideal model for addressing the relationship between these requirements and metabolic cost.

Biomechanical models of the energy expenditure of level and graded locomotion have tended to measure positive and/or negative work and assume that work is performed with mechanical efficiency values consistent with shortening and lengthening muscle contractions [1–4]. Alternatively, collisional losses of energy between the absorptive and generative phases of stance have been used to estimate the energy needed to be replaced by muscular work [5,6]. Both of these approaches do a reasonable job of explaining the relationship between changes in mechanical work and metabolic cost during different tasks. For instance, Dewolf and Willems [5] and Minetti *et al.* [7] were able to explain the optimum surface gradient for minimising the metabolic cost per distance travelled (cost of transport) of running and walking, respectively. However, the assumptions that these ‘cost of work’ models make about the muscle efficiency of generating work, along with the energy that is transferred between body segments and stored and returned elastically from tendinous tissue, leave room for this framework to be more explanatory in terms of predicting muscle behaviour and absolute changes in metabolic cost.

The cost of transport of uphill walking increases with grade and becomes a linear relationship at grades exceeding +15%; whereas for downhill walking, an increase in negative grade sees a reduction in the cost of transport, but then an increase in cost at grades steeper than −10%, with a shallower linear relationship at grades steeper than −15% [2,4]. The linear relationships at sufficiently steep uphill and downhill grades is said to equate to mechanical efficiencies of pure shortening and lengthening muscle contractions, respectively. Whereas in the shallower, non-linear region of the gradient-cost curve there may be an effect of energy-saving mechanisms, spring- and pendular-like [4,8]. The energy cost minimum at −10% grade is considered to be where mechanical energy exchange is greatest and the difference between the energy generated and energy absorbed during collisions is equal to zero [5,9]. This construct allows us to explain relationships between mechanical work and metabolic cost at a system level. However, it does not explicitly tell us what is happening to the mechanical demands of muscle across different conditions, which directly relates to energy expenditure. In particular, energy is expended for the contractile element of muscle to generate mechanical work (*w*_CE_), however inefficiencies in conversion of energy results in additional heat (*h*_CE_) produced by muscle [10], which depends on the mechanical conditions experienced and therefore influences the overall metabolic demand.

The main mechanical determinants of muscle energy use during contraction are reasonably well understood [11]. Even during isometric contraction, *h*_CE_ is produced at a given rate due to muscle activation (via ion transport), termed “activation” heat (*h*_A_), and to maintain force production (via cross-bridge cycling), termed “maintenance” heat (*h*_M_) [12–14]. These heats scale based on muscle activation level, which is proportional to the force requirements of muscle. *h*_CE_ is also produced when muscle shortens and lengthens whilst active, termed “shortening / lengthening” heat (*h*_SL_) [10,15–17]. The rate of *h*_CE_ will increase as shortening velocity increases at any given activation level relative to that of an isometric contraction (where there is no *h*_SL_). The rate of *h*_CE_ production during lengthening contractions is more complicated, initially decreasing with increased lengthening velocity, but then increasing in proportion to the work done on the muscle to stretch it [18]. *w*_CE_, which can be defined as the integral of force multiplied by the change in length of muscle and is positive during shortening and negative during lengthening [19], combines with *h*_SL_ to equal the work-related costs of muscle.

The activation and work demands of muscle are difficult to infer during locomotion because of the interactions between muscle fascicles and elastic tendinous tissues. For example, Lichtwark and Wilson [8] found that peak electromyography (EMG) of medial gastrocnemius (GM) increased with positive grade during walking between grades of +/− 10%, but interestingly with little to no change in GM fascicle shortening or shortening velocity due to the substantial elastic contributions of the Achilles tendon. Based on trends in their data (see Fig. 3A from Lichtwark and Wilson [8]) together with the linear regions of the gradient-cost curve, we might expect to see an increase in fascicle length changes at steeper uphill gradients.

Few studies have attempted to assess the activation or work demands of muscle during up- or down-hill locomotion. One of the key muscle-level indices to examine is relative activation, which likely signifies changes in both *h*_A_ and *h*_M_, as it indicates the number of fibres that need to be activated to generate force. Lay *et al.* [20] and Franz and Kram [21] found that during walking between grades of +/− 15% the mean EMG of GM, soleus (SOL), vastus medialis (VM), rectus femoris (RF), biceps femoris (BF) and gluteus maximus (GMAX) increased with incline walking, and that of VM and RF also increased with decline walking. Lay *et al.* [20] also found that these changes in muscle activity were consistent with changes in joint moments. Thus, costs of activating muscle and generating force may increase at steeper inclines and declines of walking. Meanwhile, others have measured joint work during up- and down-hill locomotion, as an indication of whether muscles act to generate or dissipate net mechanical energy (*w*_CE_). These studies show that the net mechanical energy injected by the ankle, knee and hip all increase with incline walking [22–24]. Thus, muscle work-related costs may increase as grade becomes more positive, considering also the excess heat involved in shortening muscle to generate work.

Computational musculoskeletal modelling methods [25–27] provide new avenues to explore how movement influences muscle states (e.g. activation, fibre velocity) and subsequent energetic cost [28]. Koelewijn *et al.* [29] found very strong correlations between the energy expenditure computed from several of these cost models and that calculated from indirect calorimetry on walking trials with different inclines (−8%, 0%, +8%). One of the models that was tested, that was first developed by Umberger *et al.* [28] and has since been adapted [30,31], has been evaluated against direct measures of muscle energy consumption from isolated muscle preparations, and uses muscle states to derive *ḣ*_A_, *ḣ*_M_, *ḣ*_SL_, *ẇ*_CE_, and thus the total rate of energy expenditure. Moreover, these individual rates can be partitioned from the total rate of energy expenditure that the model yields. However, the accuracy of this modelling is dependent on the accuracy of the underlying simulated muscle states.

Therefore, we aimed to assess simulated muscle mechanics and energetics from a musculoskeletal modelling approach involving the Umberger metabolic cost model, and then to use this approach to estimate how muscle force- (*ḣ*_A_ + *ḣ*_M_) and muscle work-related cost (*ẇ*_CE_ + *ḣ*_SL_) influence the total rate of energy expenditure across different uphill and downhill walking conditions (changes to mechanical work requirements) between +/− 15% grade. Note, we define muscle force-related costs as the costs associated with activating muscle to achieve force requirements. Therefore, they are also linked to muscle dynamics (i.e. when force potential drops there is a greater requirement for activation to meet the force requirements). We hypothesised that muscle force-related costs would rise at steeper inclines (i.e. steeper positive grades) and steeper declines (i.e. steeper negative grades), and that muscle work-related costs would predominantly rise as grade became more positive; and therefore that changes in force-related costs would dominate the changes in total rate of energy expenditure during downhill walking, with more of a contribution of work-related costs during uphill walking. We also expected that muscle force-related costs and muscle work-related costs would be at least moderately correlated to joint moments and joint work, respectively.

## Materials and methods

### Participation

Thirteen healthy participants (5 female, 8 male), age 30 ± 7 (mean ± SD) years, height 179 ± 9 cm, and mass 76 ± 10 kg consented to participate in this study. This study was approved by, and conducted in accordance with the University of Queensland, Faculty of Health and Behavioural Sciences Low and Negligible Risk Ethics Sub-Committee (2022/HE000001). This study conformed to the standards set by the Declaration of Helsinki.

### Experimental protocol

Participants walked on a force-instrumented tandem-belt treadmill (Tandem Treadmill; AMTI, MA, USA). Treadmill speed was set to each participant’s preferred walking speed (1.2 ± 0.1 m·s^−1^). Participants selected their preferred walking speed during a 3 min familiarisation period that involved varying the treadmill speed between 0.8 to 1.5 m·s^−1^ on level gradient. Seven different fixed gradient conditions – three downhill (−15%, −10%, −5%), one level (0%) and three uphill (+5%, +10%, +15%) – were performed at random during a single testing session. Each condition lasted 5 min, and each was separated by a minimum of 5 min rest, until subjects felt fully recovered. Metabolic energy consumption was recorded throughout each trial and kinematics, kinetics and muscle mechanics were recorded simultaneously for 10 sec at the 3 min mark of each trial; under the assumption that steady-state walking was reached by the 3 min mark.

### Experimental data

#### Kinetics and kinematics

Ground reaction force (GRF) data was collected at 2000 Hz from the force-instrumented tandem-belt treadmill. An eight-camera motion capture system (Vantage, Vicon, Oxford, UK) was used to capture the position of 43 reflective markers at 200 Hz and was time-synchronised with the GRF data. Markers were placed on anatomical landmarks on both feet, shanks and thighs and on the pelvis and torso [32]. Raw marker trajectories were filtered using a zero lag fourth order low-pass Butterworth filter with a 10 Hz cut-off frequency. GRF data were filtered using a zero lag second order low-pass Butterworth filter with a 10 Hz cut-off frequency, which was needed to filter out treadmill noise and improve force assignment. Marker and force data were then digitally exported to OpenSim [26] for analysis. An OpenSim model developed by Lai *et al.* [33] was scaled using OpenSim’s Scale Tool. Joint kinematics during walking were then determined using OpenSim’s inverse kinematics tool. Joint kinematics and GRF data were then passed through OpenSim’s inverse dynamics tool, where kinematics were filtered at 15Hz with zero phase shift using OpenSim’s built-in filter, and joint kinetics were computed. The joint kinematics and kinetics were then imported into MATLAB for further analysis (MathWorks, MA, USA). A custom MATLAB script was used to differentiate joint angles with respect to time to determine joint velocities, which were then multiplied with joint moments to determine joint powers. Time-series data were cropped from the start to the end of the gait cycle (i.e. from right-foot ground contact through to the end of the subsequent right-foot swing phase; events indexed based on GRF data). Stride metrics (duty factor and ground contact time) are detailed below (Table 1). Joint moments were averaged over the gait cycle to represent the mean force requirements across each joint. Joint powers were integrated with respect to time over the gait cycle to determine the net work requirements of each joint. All kinetic and kinematic data, along with the following muscle fascicle and muscle activation data, from each condition that each participant performed was averaged over three gait cycles.

**Table 1.** The mean ± SD gait cycle duration (t_cycle_) and duty factor across all prescribed walking conditions (−15%, −10%, −5%, 0%, +5%, +10%, +15%). Data averaged across twelve participants.

|  | -15% | -10% | -5% | 0% | +5% | +10% | +15% |
| --- | --- | --- | --- | --- | --- | --- | --- |
| t<sub>cycle</sub> (s) | 1.14 ± 0.10 | 1.14 ± 0.08 | 1.15 ± 0.07 | 1.17 ± 0.06 | 1.20 ± 0.08 | 1.23 ± 0.10 | 1.20 ± 0.12 |
| duty factor | 0.64 ± 0.01 | 0.64 ± 0.01 | 0.65 ± 0.01 | 0.65 ± 0.01 | 0.65 ± 0.01 | 0.65 ± 0.01 | 0.66 ± 0.01 |

#### Muscle fascicle dynamics

B-mode ultrasound (ArtUs EXT-1H, Telemed, Vilnius, Lithuania) was used to image muscle fascicle lengths in-vivo. One linear array transducer (LV8-5N60-A2, Telemed, Vilnius, Lithuania) was used to visualise GL and SOL muscle fascicles, and a second transducer of the same type was used to visualise VL muscle fascicles. Both transducers were fixed to the participant using self-adhesive bandage (Medichill, WA, Australia) such that they maintained longitudinal alignment to the plane of the fascicles during walking [34]. Probe depth was set to 50 mm and images were recorded using EchoWave II software (Telemed, Vilnius, Lithuania) which was operated on two separate laptop computers. A frame-by-frame external trigger was used to collect data from both transducers at 160 Hz and to synchronise this data to the motion capture and GRF data. Data from the transducers were exported to MATLAB where fascicle tracking software [34,35] was used to determine fascicle length measurements. This software performed automated tracking of a manually defined fascicle using an affine flow algorithm. Manual corrections were made if realignment of a measured fascicle was necessary. The same fascicle region was tracked across all trials for each participant. Fascicle lengths were cropped from the start to the end of the gait cycle. Each participant’s muscle fascicle lengths were normalised to the respective muscle’s mean fascicle length across all conditions that the participant performed.

#### Muscle activation

Surface EMG was used to measure the activation of GL, GM, SOL, tibialis anterior (TA), VL, RF, BF and GMAX. Recording sites were prepared by shaving and then cleaning the shaved area using an abrasive gel (NuPrep, Weaver and Company, CO, USA) and alcohol. 24 mm diameter EMG electrodes (Covidien, MA, USA) were placed in a bipolar configuration over each muscle, according to SENIAM guidelines [36]. EMG signals were acquired using wireless sensors (MiniWave, Cometa, MO, USA) and were amplified by 1000-times, hardware filtered with a 10 Hz high-pass filter and a 1000 Hz low-pass filter (WavePlus, Cometa, MO, USA) and recorded in Vicon Nexus at 2000 Hz. EMG signals were then processed in MATLAB where DC offsets were removed, a 25 Hz high-pass filter was applied, followed by signal rectification, and then a zero-lag second order Butterworth filter with a 10 Hz cut-off frequency was applied forward and back in time, producing an envelope [37]. The filtered EMG signals were then cropped from the start to the end of the gait cycle. Each EMG signal from each participant was normalised to the respective muscle’s maximum enveloped EMG across all conditions that the participant performed.

#### Metabolic energy consumption

Oxygen consumption and carbon dioxide expiration were recorded using a portable spirometry system (Metamax 3B; Cortex, Leipzig, Germany). Testing was performed 2 hr post-prandial and prior to any daily caffeine consumption to mitigate the effects of diet on metabolic rate. Resting values were obtained over a 5 min period of quiet standing prior to beginning the trials. To ensure primarily oxidative metabolism, trials were terminated where the respiratory exchange ratio (RER) rose above 1.0. The rate of oxygen consumption was required to reach a plateau (i.e. steady-state respiration) within the first 3 min of each condition. This plateau was confirmed visually and by performing regression analyses of the final 2 min of oxygen consumption data of each trial to ensure high linearity (r^2^ > 0.95). The data from the final 2 min of each trial was averaged and converted to gross metabolic power using standard equations to obtain a representative steady-state value [38].

### Neuromuscular simulations

We used the open-source model published by Lai *et al.* [33] as the base model for our analyses. This model was comprised of 80 Hill-type muscle-tendon units. Nineteen degrees of freedom (6 at the pelvis, 5 in each leg, and 3 in the upper body) were considered in the model. We found this model to be more suitable than others because of its design to reduce passive forces when simulating movements that involve substantial hip and knee flexion (e.g. steep uphill walking). To verify that the experimental motion capture data was being correctly applied to the model, joint kinematics and kinetics were compared visually to graded walking data reported in the literature. We also ensured that the reserve moments (the difference between required net joint moments and muscle-generated net joint moments) were no greater than 5% of the peak net joint moments, as per recommendations [39,40].

To solve the redundancy problem and determine muscle activations to achieve the required dynamics, we used OpenSim Moco (v1.2.0) software [39]. OpenSim Moco uses direct collocation to solve optimal control problems; using body kinematics and kinetics that we prescribed and a cost function that minimises the sum of square controls to solve for muscle excitations and fascicle length changes. We first used an automated approach to optimise the muscle properties of each participant’s model [41]; scaled muscles’ maximum isometric force based on the mass of the participant relative to the mass of the generic model; adjusted models’ GL, SOL, and VL optimal fascicle lengths to the respective mean fascicle lengths to which we normalised our experimental data; adjusted the tendon slack lengths to account for these changes in optimal fascicle length (to maintain a similar joint angle at which muscle fascicle length first starts to increase when the joint is passively rotated); and converted the muscle model type to *DeGrooteFregly2016Muscle* [42]. We then went through a systematic process to determine whether any model properties could be manually adjusted to reduce absolute differences and improve the qualitative similarity of salient features between our experimental data and simulated muscle activations (Fig. 1), fascicle length changes (Fig. 2) and whole-body metabolic powers (Fig. 3). This “evaluation” of the modelling was performed on seven of our thirteen subjects who varied in mass and sex (61 - 92 kg; 3 female, 4 male). We found that the outputs from the simulation were highly consistent between subjects so we were confident that adding more subjects to this evaluation would not affect the results from it.

**Figure 1.**
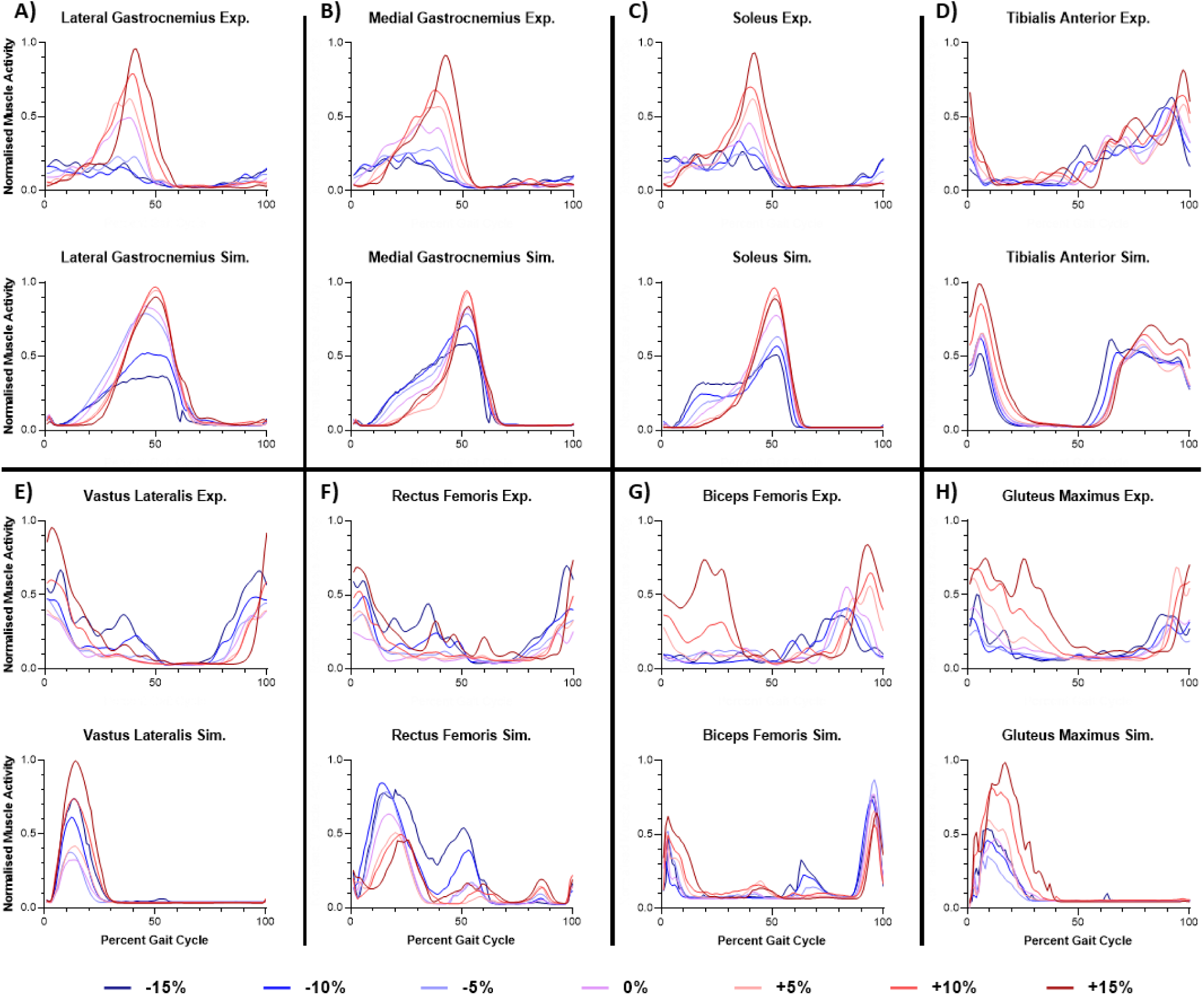
Comparison between normalised experimental EMG and normalised simulated muscle activity across the gait cycle (time-normalised from right-foot ground contact through to the end of the right-foot swing phase) and across all prescribed walking conditions (−15%, −10%, −5%, 0%, +5%, +10%, +15%; dark blue to dark red, respectively) for (**A**) GL, (**B**) GM, (**C**) SOL, (**D**) TA, (**E**) VL, (**F**) RF, (**G**) BF and (**H**) GMAX. Data averaged across seven participants.

**Figure 2.**
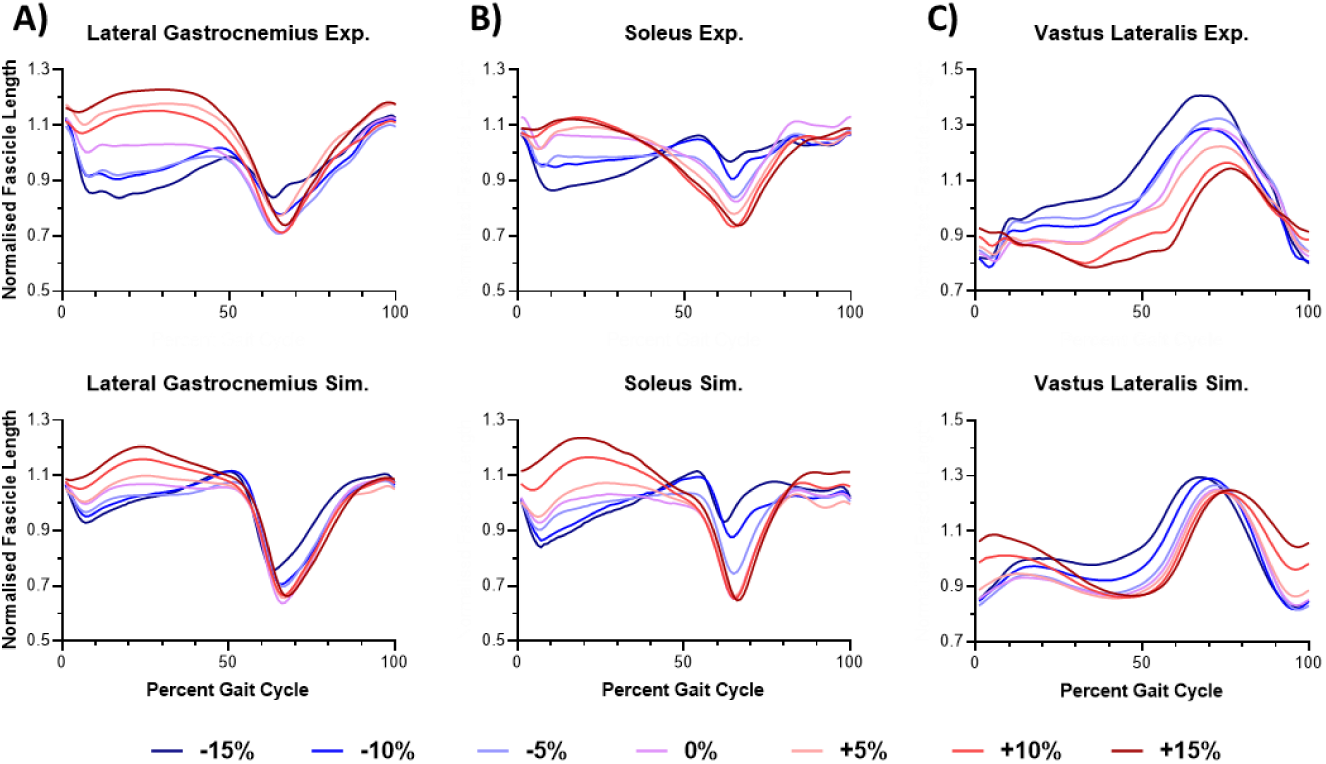
Comparison between normalised experimental and simulated fascicle length changes across the gait cycle (time-normalised from right-foot ground contact through to the end of the right-foot swing phase) and across all prescribed walking conditions (−15%, −10%, −5%, 0%, +5%, +10%, +15%) for (**A**) GL, (**B**) SOL and VL (**C**). Data averaged across seven participants.

**Figure 3.**
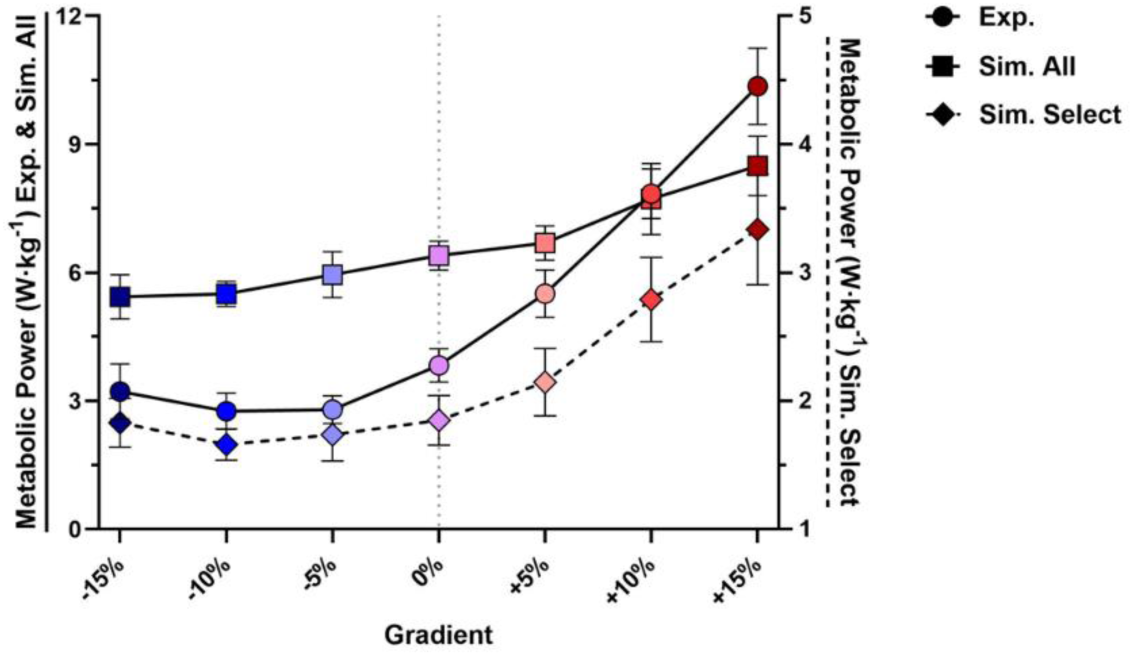
Comparison between mean ± SD experimental net metabolic power (W·kg^−1^) (*Exp.*; circles; solid lines; primary Y-axis), simulated net metabolic power summed from all muscles in the musculoskeletal model (*Sim. All*; squares; solid lines; primary Y-axis) and simulated net metabolic power summed from only GL, GM, SOL, TA, VL, RF, BF and GMAX in the musculoskeletal model (*Sim. Select*; diamonds; dashed lines; secondary Y-axis) across each of the prescribed walking conditions (−15%, −10%, −5%, 0%, +5%, +10%, +15%). *Exp.* and *Sim. All* data are averaged across seven participants, while *Sim. Select* data are averaged across twelve participants.

Due to the significant computational time that it takes to simulate gait cycles in Moco, we elected to simulate a single gait cycle per trial, per participant. The particular gait cycle that we chose to simulate was based visually on 1) the symmetry between left and right ankle, knee and hip moments, and 2) the magnitude and shape of these moments compared to the average of the three gait cycles across which the experimental data were averaged. Simulated muscle activations and fascicle length changes were collated for the same muscles measured experimentally for the evaluation. Each of these simulated muscle activations from each of the seven subjects was normalised to the respective muscle’s maximum simulated activation across all trials that the participant performed, as per the process of normalising the experimental EMG. Simulated fascicle lengths were normalised to the same mean values as the experimental data.

Whole-body metabolic power was estimated from the simulations performed on those seven subjects using the model developed by Umberger *et al.* [28] with some modifications [30,31]. The Umberger model was implemented via OpenSim’s probe reporter tool using the *Umberger2010MuscleMetabolicsProbe*. To compute gross whole-body metabolic power, the controls and states from Moco were fed to the cost model, we then summed the computed rate of energy expenditure across all muscles, plus the outputted whole-body mass-specific basal rate, and then integrated the resulting whole-body metabolic rate over the gait cycle and divided by the duration of the gait cycle.

During the process of model refinement within our evaluation to achieve suitable muscle activations and muscle fibre trajectories, we found it necessary to remove the gastrocnemii femoral and tibial wrapping surfaces from the model because they led to overactivation and discontinuities in fascicle length changes of the gastrocnemii. We also locked the subtalar and metatarsophalangeal joints in the model to minimise instability that otherwise caused overactivation of the muscles that spanned those joints. Further, we added reserve actuators to each model degree-of-freedom with a maximum force of 10.0 N or Nm to combat any remaining dynamic inconsistency in the modelling. Additionally, after running simulations across all thirteen participants with the validated model, we excluded one male participant (age 18 years, height 196 cm, and mass 83.6 kg), who was not one of the seven participants in the aforementioned evaluation, from the group analysis due to an overactivation of muscles spanning the hip and knee that we were unable to reduce or resolve.

#### Metabolic energy breakdown

With our modelling approach evaluated and musculoskeletal simulations then performed on our twelve subjects, we used the Umberger metabolic cost model to partition the contributions of muscle force- (*ḣ*_A_ + *ḣ*_M_) and work-related (*ẇ*_CE_ + *ḣ*_SL_) costs to simulated total metabolic power across the different walking conditions. We analysed the simulated whole-body metabolic power using all muscles in the model (Fig. 3), as well as only those for which we could evaluate based on EMG and ultrasound measures (Fig. 4A). Large absolute differences between simulated and experimental metabolic power were found in the former analysis; and after examining the mean percentage of total simulated metabolic power from each muscle in the model across the graded walking conditions (Fig. S1), it seemed that muscles such as those involved in stabilising and flexing the hip joint (e.g. gluteus medius, iliopsoas) were contributing too much. In contrast, the correlation between simulated and experimental metabolic power was considerably stronger in the latter analysis, and thus we decided to factor only those muscles that we had evaluated (GL, GM, SOL, TA, VL, RF, BF and GMAX) into our analyses of muscle activation- and work-related costs.

**Figure 4.**
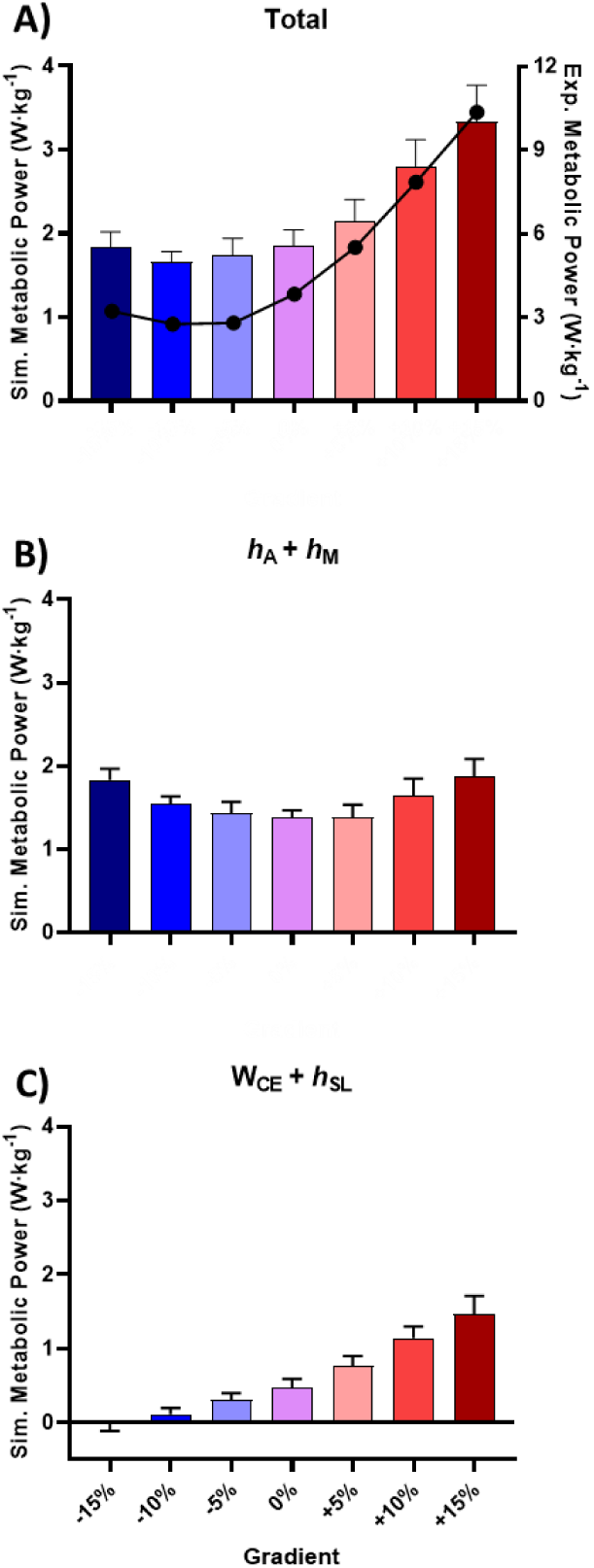
Partitioning mean ± SD simulated net metabolic power (W·kg^−1^) summed from GL, GM, SOL, TA, VL, RF, BF and GMAX into force- and activation-related costs across all prescribed walking conditions. (**A**) Total metabolic power. (**B**) Force-related costs (activation heat (ḣ_A_) + maintenance heat (ḣ_M_)). (**C**) Work- related costs (work of contractile element (ẇ_CE_) + shortening/lengthening heat (ḣ_SL_)). (**A**) Also includes the mean experimental metabolic power (W·kg^−1^) for each walking condition (circles) on a separate axis. Simulated data are averaged across twelve participants, while experimental data are averaged across seven participants.

#### Statistics

We tested whether simulated metabolic power summed across only GL, GM, SOL, TA, VL, RF, BF and GMAX was more strongly correlated to our experimental metabolic power across the graded walking conditions, compared to simulated metabolic power summed across all muscles in the model. We used linear models (LMs) and accounted for each participant as a categorical predictor to account for variation due to participant in the data used in the regression. The resulting adjusted correlation coefficient (R^2^) between each form of simulated metabolic power and the experimental metabolic power was used to determine their strength of association.

We also used LMs to determine the strength of association between muscle activation- and work-related costs and mean joint moment and net joint work, respectively. We ran LMs for ankle, knee and hip mean joint moment and net joint work to determine how well each of the joints explained variance in muscle activation- and work-related costs across the graded walking conditions. We also ran LMs that included all of the joints to test how much this would strengthen the resulting adjusted R^2^. Again, we accounted for participant in each of the LMs. We considered R^2^ values of ≤ 0.35 as weak correlations, 0.36 to 0.69 as moderate correlations, 0.70 to 0.89 as strong correlations, and ≥ 0.90 as very strong correlations [43].

## Results

### Stride metrics

Despite having only walking speed constrained, we found that participants used a similar gait cycle duration and duty factor across all conditions (Table 1).

### Model validation

Visual assessment of the shape and magnitude of simulated muscle activations and experimental data across the prescribed walking conditions (Fig. 1) demonstrated reasonable similarities, although some differences did exist (e.g. pre-contact activation of VL and GMAX). We note that the timing of the simulated data was delayed relative to the experimental data. We observed an increase in the activation of GL, GM, SOL, TA, BF and GMAX as grade became more positive, and of VL and RF at steeper positive and negative grades.

Simulated GL, SOL and VL muscle fascicle length changes were visually similar in shape, magnitude and timing to the experimental data across the prescribed walking conditions (Fig. 2). As grade became less negative, from −15% to +15% grade, fascicles went from lengthening more to shortening more during stance.

We observed large absolute differences, yet a strong correlation (adjusted R^2^ = 0.75), between experimental and simulated metabolic power when accounting for costs from all muscles in the musculoskeletal model (Fig. 3; Fig. S2A). Simulated metabolic power summed across all muscles in the model was ~40% higher than that measured experimentally at 0% grade, which generally persisted through the downhill conditions, although simulated data did not increase as much as that measured experimentally in uphill grades.

### Metabolic energy expenditure

Despite being on a smaller scale, simulated metabolic power summed across only GL, GM, SOL, TA, VL, RF, BF and GMAX was very strongly correlated with experimental metabolic power (adjusted R^2^ = 0.91) across the graded walking conditions (Fig. 3; Fig. 4A; Fig. S2B); which is a substantial improvement compared to accounting for every muscle in the model in the simulations of metabolic power. Using only these select walking muscles, we observed a U-shaped relationship between total metabolic power and gradient between 0 to −15% grade, with the minimum at −10% grade, and a progressive increase in total metabolic power between 0 to +15% grade; which are the same trends observed in the experimental data. Muscle force costs (*ḣ*_A_ + *ḣ*_M_) increased at steeper uphill and downhill gradients (Fig. 4B), and muscle work costs (*ẇ*_CE_ + *ḣ*_SL_) increased as grade became more positive (Fig. 4C).

### Joint mechanics

There was a U-shaped relationship between knee mean moment and gradient, increasing at steeper uphill and downhill grades (Fig. 5), and there was relatively no change in ankle and hip mean moment across different grades. Knee mean moment was moderately correlated to muscle force costs (adjusted R^2^ = 0.38) and ankle and hip mean moment were weakly correlated to muscle force costs (adjusted R^2^ = 0.06 and 0.13, respectively). Including ankle, knee and hip mean moment as predictors in the regression model explained substantially more variance in muscle force costs than when considered singularly (adjusted R^2^ = 0.63) (Fig. S3A).

**Figure 5.**
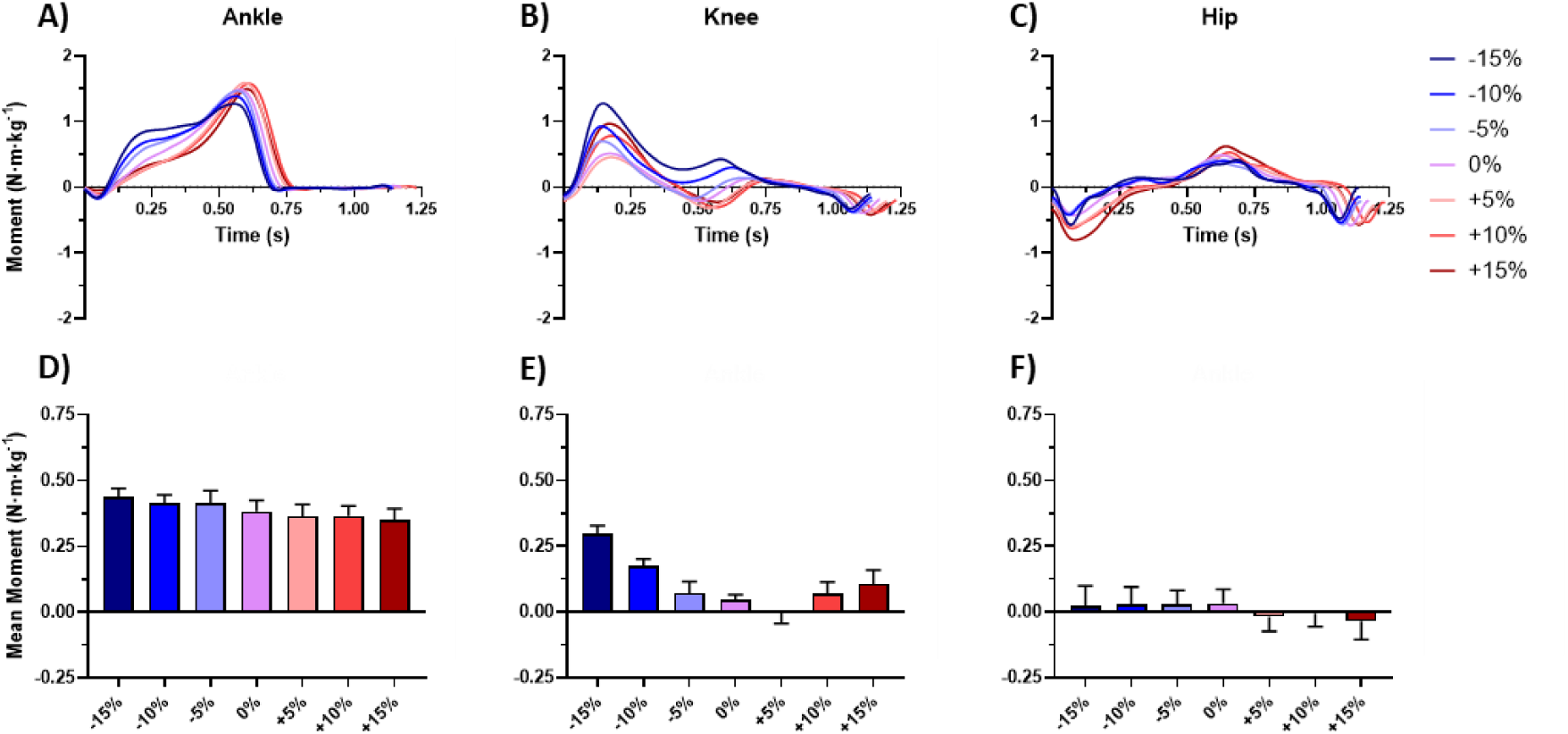
Comparison of (**A**) ankle, (**B**) knee and (**C**) hip joint moments (N·m·kg^−1^) across the gait cycle (real- time, from right-foot ground contact through to the end of the right-foot swing phase), and (**D**) ankle, (**E**) knee and (**F**) hip mean ± SD mean moment, across all prescribed walking conditions (−15%, −10%, −5%, 0%, +5%, +10%, +15%). Data averaged across twelve participants.

Across all three joints, net joint work became more positive as gradient became more positive (Fig. 6). There was also a strong correlation between ankle, knee and hip net joint work and muscle work costs (adjusted R^2^ = 0.88, 0.80 and 0.70, respectively). Moreover, including the net joint work of all three joints as predictors in the regression model explained almost all of the variance in muscle work costs (adjusted R^2^ = 0.97) (Fig. S3B). Interestingly, positive joint work, especially of the knee, increased at steeper uphill grades and plateaued during level and downhill walking, similar to total rate of energy expenditure (Fig. 4A). Including the positive joint work of all three joints in a regression model against the total rate of energy expenditure resulted in a very strong correlation (adjusted R^2^ = 0.94) (Fig. S3C).

**Figure 6.**
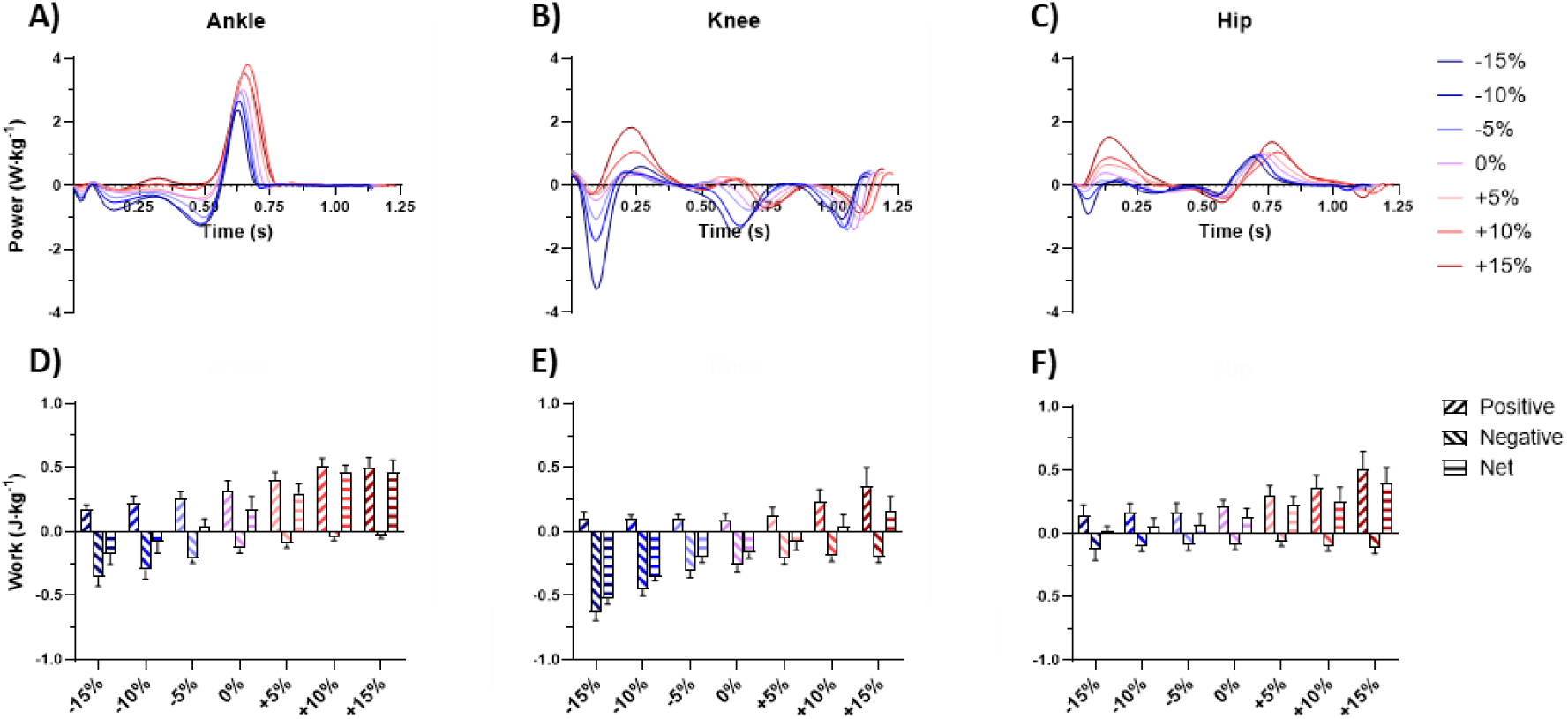
Comparison of (**A**) ankle, (**B**) knee and (**C**) hip joint powers (W·kg^−1^) across the gait cycle (real- time, from right-foot ground contact through to the end of the right-foot swing phase), and (**D**) ankle, (**E**) knee and (**F**) hip mean ± SD positive, negative and net joint work (J·kg^−1^), across all prescribed walking conditions (−15%, −10%, −5%, 0%, +5%, +10%, +15%). Data averaged across twelve participants.

## Discussion

We used a musculoskeletal modelling approach to simulate muscle mechanics and energetics across graded walking conditions between +/− 15% grade. Whilst the predictions of whole body metabolic rates were substantially different in absolute terms to the experimental data, when we factored in the rates of energy expenditure from only the major work producing muscles in the model, the simulated and experimental metabolic rates became more strongly correlated. As hypothesised, muscle force-related costs (*ḣ*_A_ + *ḣ*_M_) increased as uphill and downhill grades became steeper and muscle work-related costs (*ẇ*_CE_ + *ḣ*_SL_) increased as grade became more positive, and hence, force costs dominated the changes in metabolic rate during downhill walking with more of a contribution of work costs during uphill walking.

Muscle force costs (Fig. 4B) were predicted reasonably well by changes in mean joint moment (Fig. 5D-F; Fig. S3A). We observed a similar U-shaped relationship between force costs and grade as we did knee mean joint moment and grade, increasing at steeper up- and down-hill grades; with little to no change in ankle or hip mean joint moment with grade. This was mirrored by the contribution of different musculature to changes in total force costs across the different conditions (Fig. 7), in which VL and RF demonstrate the greatest changes across grades, among the muscles that were measured. It is interesting that this is despite clear changes in the activation of ankle and hip musculature across grades (Fig. 1), in line with the findings of Lay *et al.* [20] and Franz and Kram [21]. However, the scale of these changes in activation, mean moment and force costs are much smaller at the ankle and hip compared to the knee.

**Figure 7.**
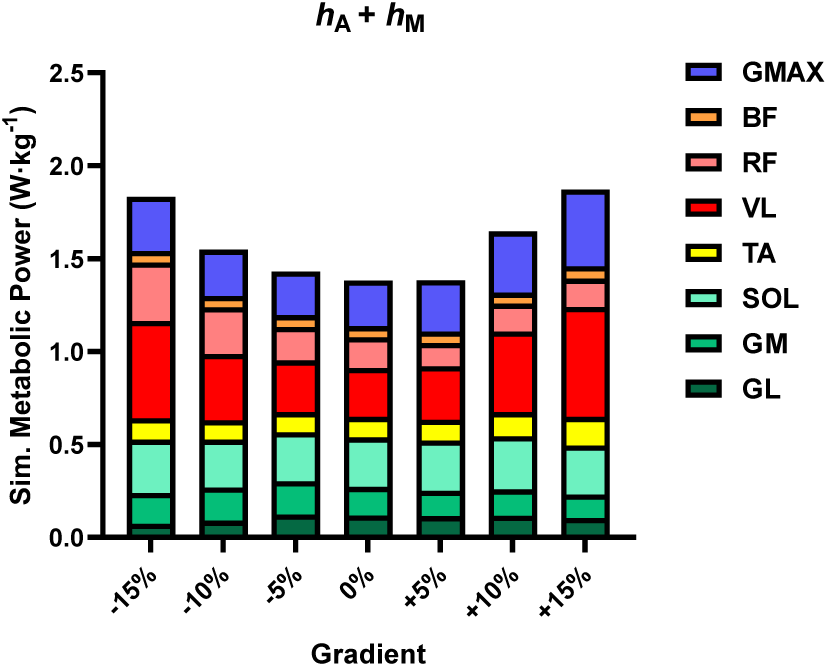
Comparison between the contribution of GL, GM, SOL, TA, VL, RF, BF and GMAX to total activation-related costs (activation heat (ḣ_A_) + maintenance heat (ḣ_M_)) (W·kg^−1^) across all prescribed walking conditions (−15%, −10%, −5%, 0%, +5%, +10%, +15%) from the Umberger cost model. Data averaged across twelve participants.

We observed an increase in muscle work costs as grade became more positive (Fig. 4C), which was predicted extremely well by changes in net joint work across the ankle, knee and hip (Fig. 6D-F; Fig. S3B). Based on net joint work, we assume that ankle, knee and hip musculature each contributed substantially to changes in total work costs. Unsurprisingly, as grade became more positive – rising from −15% to +15% grade – GL, SOL and VL fascicles went from lengthening more to shortening more during stance (Fig. 2). Furthermore, the total metabolic rate was very strongly correlated with changes in positive joint work (Fig. S3C). This is pertinent because collisional models tend to consider only positive work as costly (assuming that the costs of isometric and eccentric contractions are relatively negligible) and, thus, base their calculations on the positive work needed to make up for energy dissipation during stance [6]. It is interesting that, similar to total metabolic rate, we observed a plateau in positive joint work as grade became more negative from 0% to −15% grade (Fig. 6D-F); which we suspect is a result of the increase in force requirements as downhill grade became steeper. However, there are some slight dissimilarities between the changes in positive joint work and total metabolic rate, such as where the energetic minimum lies, along with the uptick in total metabolic rate at steeper downhill conditions.

The gradient-cost curve of walking is characterised by a U-shaped relationship between 0 to −15% grade, with the minimum at −10% grade, and a progressive increase in cost between 0 to +15% grade. Muscle force costs made up the majority of the total rate of energy expenditure across all conditions, particularly downhill, and contributed to the non-linearity of the gradient-cost curve; while muscle work costs explained how the cost of walking uphill far exceeded that of walking downhill. It is likely that the increases in knee joint moments and force costs were driven by higher braking force requirements at steeper downhill grades and higher propulsive force requirements at steeper uphill grades [44–46]; and that the increases in work demands were driven by increases in vertical mechanical power required to raise the body against gravity [1,5,47].

Whilst joint work was able to explain almost all of the variance in muscle work costs, we found some dissociation between joint moments and muscle force costs. This is because the costs of generating force will not only scale based on the magnitude of force, but also length at which muscle is producing force, per the force-length relationship [10,48,49]. For instance, in Fig. 8, we can see that the activation of VL and force requirements of the knee are higher at −15% compared to 0% grade, however the length of VL fascicles during the main period of force production are closer to energetically optimal (i.e. closer to 1.0) at −15% compared to 0% grade. We used mean joint moments as an indicator of the average force requirements over time [50,51]. Since gait cycle duration and duty factor were unchanged across the conditions (Table 1), these average force requirements are proportional to other metrics of force and time that have been used to try to predict the cost of generating force (e.g. [52–56]). Therefore, current force-based cost models may be unable to explain some variance in muscle force costs across different muscle length conditions.

**Figure 8.**
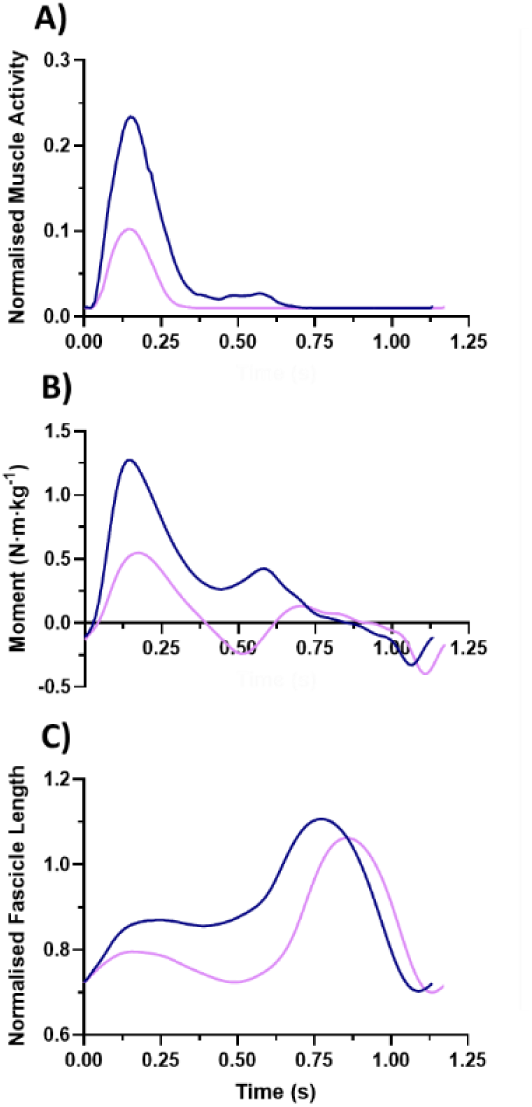
Comparison of time-series (**A**) simulated normalised muscle activity of VL, (**B**) knee joint moment (N·m·kg^−1^) and (**C**) simulated normalised fascicle length of VL between walking at −15% (blue) and 0% (purple) grade. Data averaged across twelve participants.

Our findings have important implications for biomechanical models of locomotor energetics. Other research has shown that mechanical work- and collision-based cost models (e.g. [2,3,5]) are sufficient for explaining relationships between mechanical work and metabolic cost at a system level, but likely are unable to predict other costs associated with force production. In the context of graded walking, we suspect that these models would be able to account for changes in muscle work-related costs, but that an additional cost term is needed to account for muscle force-related costs; which seems especially important for describing costs of downhill walking and non-linearity in the gradient-cost curve. Notably, Hoogkamer *et al.* [45] achieved better estimates of the energy expenditure of uphill running by including both force- and mechanical work-based cost terms in their model, as opposed to models based purely on mechanical work. Furthermore, our results suggest that models could use knee mechanics as an indicator of the energy costs of up- and down-hill walking. However, it must be noted that our participants were constrained to a single walking speed and that other gait metrics were similar across the conditions (Table 1) – whether changes to gait parameters alter the joint-level contributions to activation and work demands and alter the shape of the gradient-cost curve, needs to be investigated [21,57,58].

This is the first study to evaluate a complete three-dimensional lower-limb musculoskeletal model for simulating muscle mechanics and energetics during up- and down-hill walking. There was good agreement between salient features of experimental and simulated muscle activations (Fig. 1) and fascicle dynamics (Fig. 2) among key walking muscles (GL, GM, SOL, TA, VL, RF, BF and GMAX) with changes in grade. However, there were large absolute differences – an overestimation at all but steep uphill conditions – compared to the experimental data by factoring all muscles in the model into our simulations of metabolic power (Fig. 3). This is a stark contrast to the findings of Koelewijn *et al.* [29] who generally observed smaller absolute differences and an underestimation of energy cost, between +/− 8% grade. Although, there are many potential reasons why our results differed, including that they performed a two-dimensional analysis in the sagittal plane that involved a custom muscle model and optimisation problem on 8 muscles (as opposed to 40) in each leg and that they used different metabolic cost models. When we included only the muscles whose mechanics we evaluated against our experimental data in our simulations of metabolic power, we observed a very strong correlation between the experimental and simulated gradient-cost curve (Fig. 4A; Fig. S2B). This indicated that mechanics of other muscles in our model were likely simulated poorly, which may have been a result of the objective function or dynamic inconsistency due to measurement error, and may have been moderated by performing a two-dimensional analysis similar to Koelewijn *et al.* [29].

There are other limitations to this study related to the muscle model and metabolic cost model. The use of a Hill-type muscle model meant that we could simulate the gross behaviour of muscle-tendon units (MTUs) with a low computational burden [59]. However, this was at the expense of capturing finer structural and functional details of the MTUs that affect energy expenditure; such as muscle-specific histology [60], muscle shape changes [61] and non-uniform tendon dynamics [62]. Additionally, physiology research is yet to precisely determine the cost of eccentric work and active lengthening, which has led to this being a major point of contention between metabolic cost models (for a review, see Miller *et al.* [27]). Whilst the cost model that we used includes terms specific to eccentric work, the different treatments of eccentric work have been shown to have a major effect on model-based predictions of locomotor energetics [27,63]. Furthermore, the cost models do not account for energy associated with ATP regeneration (via glycolysis or oxidative recovery), which also liberates heat on a slower timescale than that of the contractions [28,64].

Using a musculoskeletal modelling approach, we found that separating out the muscle force- and work-related costs can help explain different features of the gradient-cost curve of walking; with force costs explaining costs of walking downhill and non-linearity in the curve, and work costs explaining how the cost of walking uphill far exceeds that of walking downhill. Whilst joint work accounted for the variance in work costs, joint moments moderately correlated with force costs, which we put down to variance in muscle operating lengths. Still, compared to traditional mechanical work-based models, the inclusion of a force-based cost term to account for force costs should lead to a more explanatory cost model that can more directly map to the mechanical demands of muscle. We also highlight knee mechanics as an especially good indication of total energy costs of graded walking, and suggest that future studies determine how the shape of the gradient cost-curve changes alongside joint- and muscle-level force and work demands with changes in gait parameters (e.g. ground contact time, stride frequency). Further, we feel that the modelling approach that we took has utility to characterise the mechanical and energetic requirements of other forms of locomotion.

## Acknowledgements

We thank Dr. Nick Bianco for his advice during piloting of this study. We also thank all participants for their time and the University of Queensland for providing facilities to conduct this study.

## Declarations

### Author contributions

L.N.J.: conceptualization, formal analysis, investigation, methodology, project administration, visualization, writing—original draft, writing—review and editing; L.A.K.: conceptualization, methodology, supervision, writing—review and editing; A.G.C.: methodology, supervision, writing—review and editing; G.A.L.: conceptualization, funding acquisition, methodology, project administration, resources, supervision, writing— review and editing.

