## Supplementary Figures S1-3 for "Dissecting the metabolic costs of up- and down-hill walking"

### Supplementary material

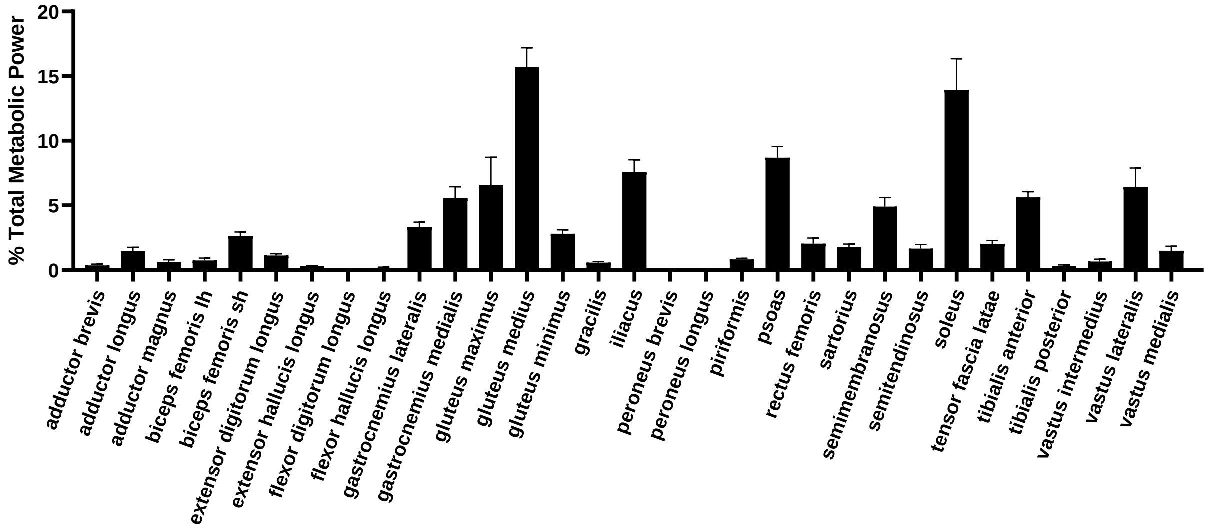


**Figure S1.** Mean ± SD contribution (%) of each muscle in the model to changes in simulated metabolic power across the graded walking conditions. Data averaged across seven participants.

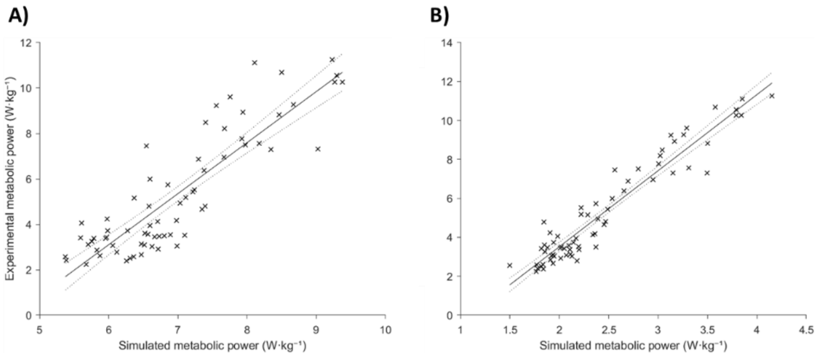


**Figure S2.** Linear models between experimental metabolic power (W·kg$^{-1}$) and (**A**) simulated metabolic power based on all muscles in the model and (**B**) simulated metabolic power based on only GL, GM, SOL, TA, VL, RF, BF and GMAX in the model. Participant was used as a categorical predictor in these linear models.

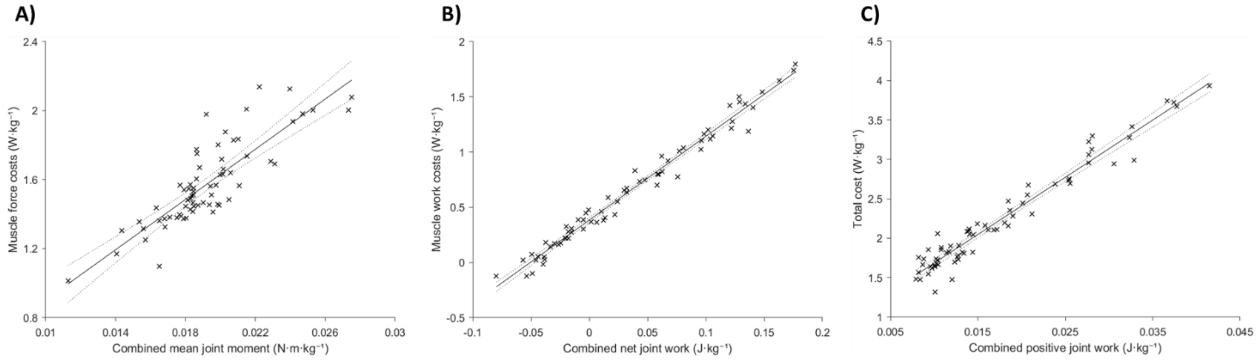


**Figure S3.** Linear models between (**A**) muscle force costs (W·kg$^{-1}$) and combined (ankle, knee and hip) mean joint moment (N·m·kg$^{-1}$), (**B**) muscle work costs (W·kg$^{-1}$) and combined net joint work (J·kg$^{-1}$) and (**C**) total cost (W·kg$^{-1}$) and combined positive joint work (J·kg$^{-1}$). Participant was used as a categorical predictor in these linear models.
